# Infant and adult brains are coupled to the dynamics of natural communication

**DOI:** 10.1101/359810

**Authors:** Elise A. Piazza, Liat Hasenfratz, Uri Hasson, Casey Lew-Williams

## Abstract

Infancy is the foundational period for learning from adults, and the dynamics of the social environment have long been proposed as central to children’s development. Here we reveal a novel, highly naturalistic approach for studying live interactions between infants and adults. Using functional near-infrared spectroscopy (fNIRS), we simultaneously and continuously measured the brains of infants (9-15 months) and an adult while they communicated and played with each other in real time. We found that time-locked neural coupling within dyads was significantly greater when they interacted with each other than with control individuals. In addition, we found that both infant and adult brains continuously tracked the moment-to-moment fluctuations of mutual gaze, infant emotion, and adult speech prosody with high temporal precision. This investigation advances what is currently known about how the brains and behaviors of infants both shape and reflect those of adults during real-life communication.

## Introduction

The ability to communicate during the first years of life requires the development of common ground with others. This involves learning to engage with our social environments using a range of behaviors, including eye gaze, facial expressions, and speech. By adulthood, we interact with others following a set of interrelated norms shared by our particular community of speakers.

Recent work with adults indicates that shared understanding is associated with shared neural responses across subjects in a set of higher-order brain regions (Hasson et al., 2012). Responses in these areas correlate strongly across subjects when incoming input is interpreted in a similar way, irrespective of its form. For example, written and spoken versions of the same story evoke comparable neural responses across readers and listeners (Regev et al., 2018). Similarly, brain responses are similar across Russophones listening to a Russian story and Anglophones listening to its English translation (Honey, Thompson, Lerner, & Hasson, 2012), and across people listening to two versions of the same story generated using synonyms (Yeshurun, Nguyen, & Hasson, 2017). Furthermore, the responses in these brain areas to the exact same ambiguous story (e.g., a short story by J.D. Salinger, or a minimalist animation) consistently differ between groups of subjects based on each group’s distinct interpretation of the story (Yeshurun et al., 2017; Nguyen, Vanderwal, & Hasson, 2018). Finally, shared neural responses in these higher-order areas were found during the successful transfer of information between a speaker’s and a listener’s brain (Stephens, Silbert, & Hasson, 2010; Zadbood et al., 2017). In these studies, the responses in the listener’s linguistic and higher-order brain areas were found to be coupled (i.e correlated with a short temporal lag) to the responses in the speaker’s brain. Furthermore, stronger speaker-listener coupling was associated with better comprehension. Speaker-listener neural coupling has been found using fMRI (Stephens et al., 2010) and functional near-infrared spectroscopy (fNIRS; Jiang et al., 2012; Liu et al., 2017), showing strong consistency between the two methodologies.

How does such a shared neural code emerge in our brains as we begin learning during the first years of life? In language, for example, the ability to understand incoming input does not emerge automatically; on the contrary, children’s understanding of the proper use of words for communicating ideas, desires, and intentions must emerge over time from ongoing interactions with other members of a community of speakers (Vygotsky, 1978; Tomasello, 1992). The child’s task is to become plugged into this community by learning from caregivers about the contextually appropriate uses of communicative signals. This generates an interesting prediction: that the ability of an infant brain to be dynamically coupled with adults’ brains may serve as a necessary precursor for early learning.

In this study, we developed a new experimental technique for simultaneously measuring an infant’s and adult’s brain during continuous, two-way interaction. Our hyperscanning paradigm opens new experimental possibilities for studying dynamic, natural communication. This approach was not available in the studies described above, which only measured neural coupling between an adult speaker and adult listeners during one-way communication using a single scanner.

Two-way interaction allows for continuous feedback from other interlocutors, through eye contact, facial expressions, and vocal prosody, among other cues. These cues may be necessary for assimilating infants into the shared language community (Tomasello, 1992). Indeed, social interactionist models emphasize the importance of social immersion for early learning (Vygotsky, 1978), and parental input is particularly instructive early in life (Brazelton, Koslowski, & Main, 1974). For example, infants use eye contact and gaze direction to infer intention, meaning, and causality (Brooks & Meltzoff, 2008; Csibra & Gergely, 2009; Senju & Csibra, 2008), and gaze synchronization is thought to enhance social connectedness between infants and adults (Brooks & Meltzoff, 2014). Infants also benefit from the rhythmic simplicity and higher, more variable pitch of infant-directed speech (Fernald & Simon, 1984) when segmenting words (Thiessen, Hill, & Saffran, 2005) and learning their meanings (Graf Estes & Hurley, 2013). Moreover, mothers modify the quality of their voices when talking to infants (Piazza, Iordan, & Lew-Williams, 2017), and change their prosody in real time based on emotional feedback from their infants (Smith & Trainor, 2008). Natural play is a key context for examining the beginnings of human cognition (Singer, Golinkoff, & Hirsh-Pasek, 2006; Fisher, Hirsh-Pasek, Newcombe, & Golinkoff, 2013).

In the present study, we aimed to understand how the brains of infants and adults become coupled to each other and to natural social behaviors in real time. To do so, we used dual-brain fNIRS to simultaneously measure the brains of an adult caregiver and 9-to-15-month-old infants while they engaged in everyday interactions, including playing, singing, and reading. fNIRS provides a non-invasive measure of changes in blood oxygenation due to neural activity, while being minimally sensitive to motion artifacts, and allows multiple participants to interact freely, face-to-face, while wearing comfortable caps (Boas, Elwell, Ferrari, & Taga, 2014). We predicted that infant-to-caregiver brain-to-brain (B2B) coupling would be higher when the infant and adult interacted with each other, relative to a control condition in which each member of the dyad interacted with another person in the room. Furthermore, we predicted that areas of the brain involved in mutual understanding (in particular, the prefrontal cortex) would track prelinguistic communicative behaviors, such as eye contact, smiling, and speech prosody.

## Results

The brain responses of the infant and the adult experimenter were recorded simultaneously using fNIRS in two conditions: *Together* and *Apart*. In the *Together* condition, each adult-infant dyad participated in face-to-face sets of playful interactions: playing an interactive game, singing a song, and reading a book. In the *Apart* condition, the neural activity of both the adult experimenter and the infant was recorded simultaneously while they interacted with other adults (i.e the infant played with a parent, and the adult conversed with another experimenter in the room). The same adult experimenter participated in interactions with all infant participants, and the order of conditions was counterbalanced (see Methods for details).

In the first set of analyses (Figures 2 and 3), we assessed the significance of neural coupling between the infant and adult experimenter in the *Together* and *Apart* conditions using a robust bootstrapped phase-scrambling analysis across all available cortical channels and a form of cross-correlation that validated the temporal specificity of coupling in one cortical ROI (the PFC). In the next set of analyses (Figures 4 and 5), we measured the relationship between the neural activation of each participant (adult, infant) and socially salient, dynamic behaviors measured throughout the *Together* condition (mutual gaze, infant smiling, and adult speech prosody). Finally, we performed control analyses to ensure that any brain-behavior correlations could not be explained by coarse task structure.

**Figure 1.**
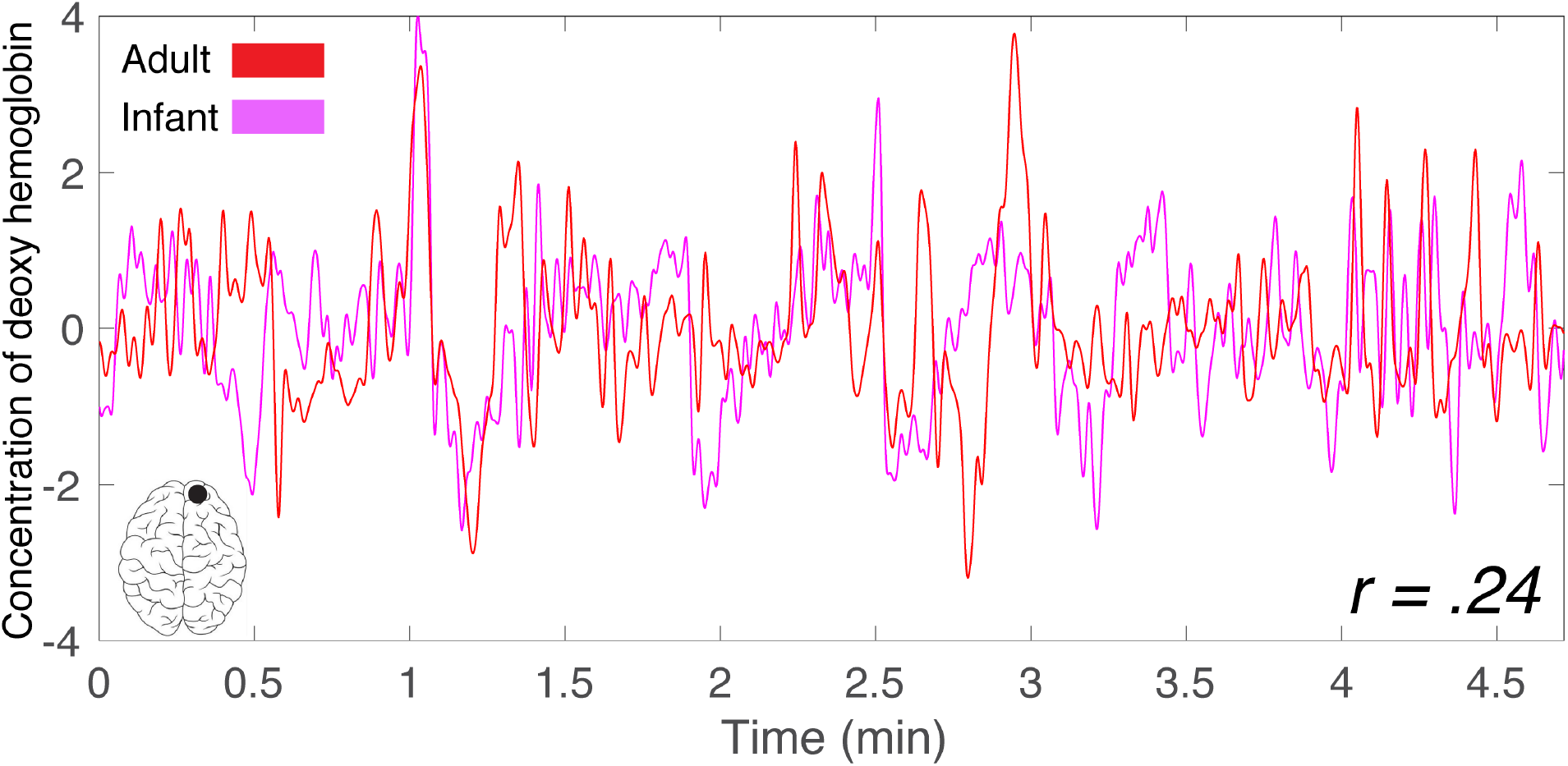
Example of the inter-subject correlation (ISC) measure, computed from a single right PFC channel of an adult (red) and an infant (pink).

**Figure 2.**
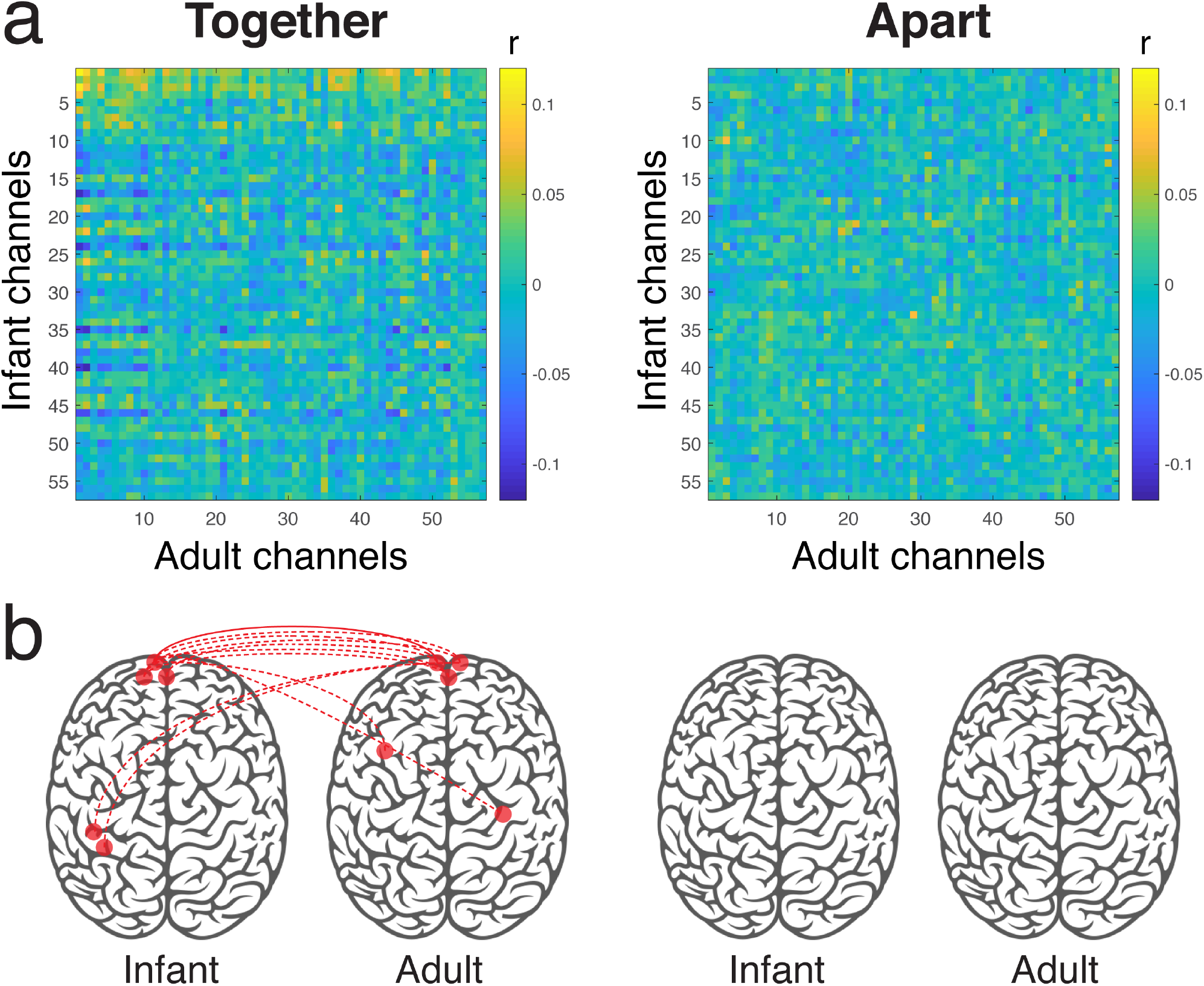
(a) Inter-subject correlation (ISC) matrices between the infant and adult. (b) Significantly coupled channel pairs, determined by phase-scrambling analysis (see Results) and corrected for multiple comparisons using false discovery rate (FDR, *q* < .05; Benjamini & Hochberg, 1995). Solid lines indicate homologous channel pairs; dotted lines indicated non-homologous channel pairs. *N* = 18 dyads.

**Figure 3.**
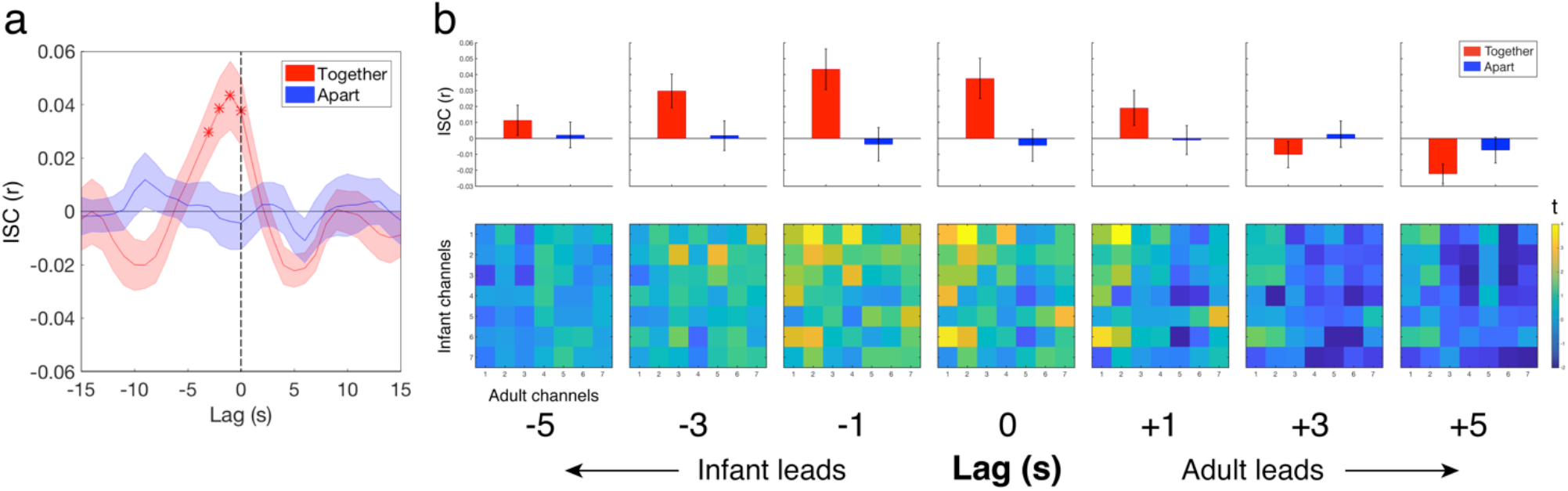
(a) Inter-subject correlation (ISC) between the adult and infant (averaged across PFC channels) in the *Together* (red) and *Apart* (blue) conditions, across a range of relative shifts of the two signals. ISC was stronger in the *Together* than the *Apart* condition when the adult’s and infant’s signals were temporally aligned (no lag) or shifted relative to each other by up to -3 s (i.e infant leading adult). Stars indicate significant lags, corrected for multiple comparisons. (b) Top: mean ISC values across all 49 pairwise (adult-infant) combinations of PFC channels, averaged across dyads. Bottom: *t*-map comparing ISCs (*Together* vs. *Apart*) for each PFC channel combination. *N* = 18 dyads.

**Figure 4.**
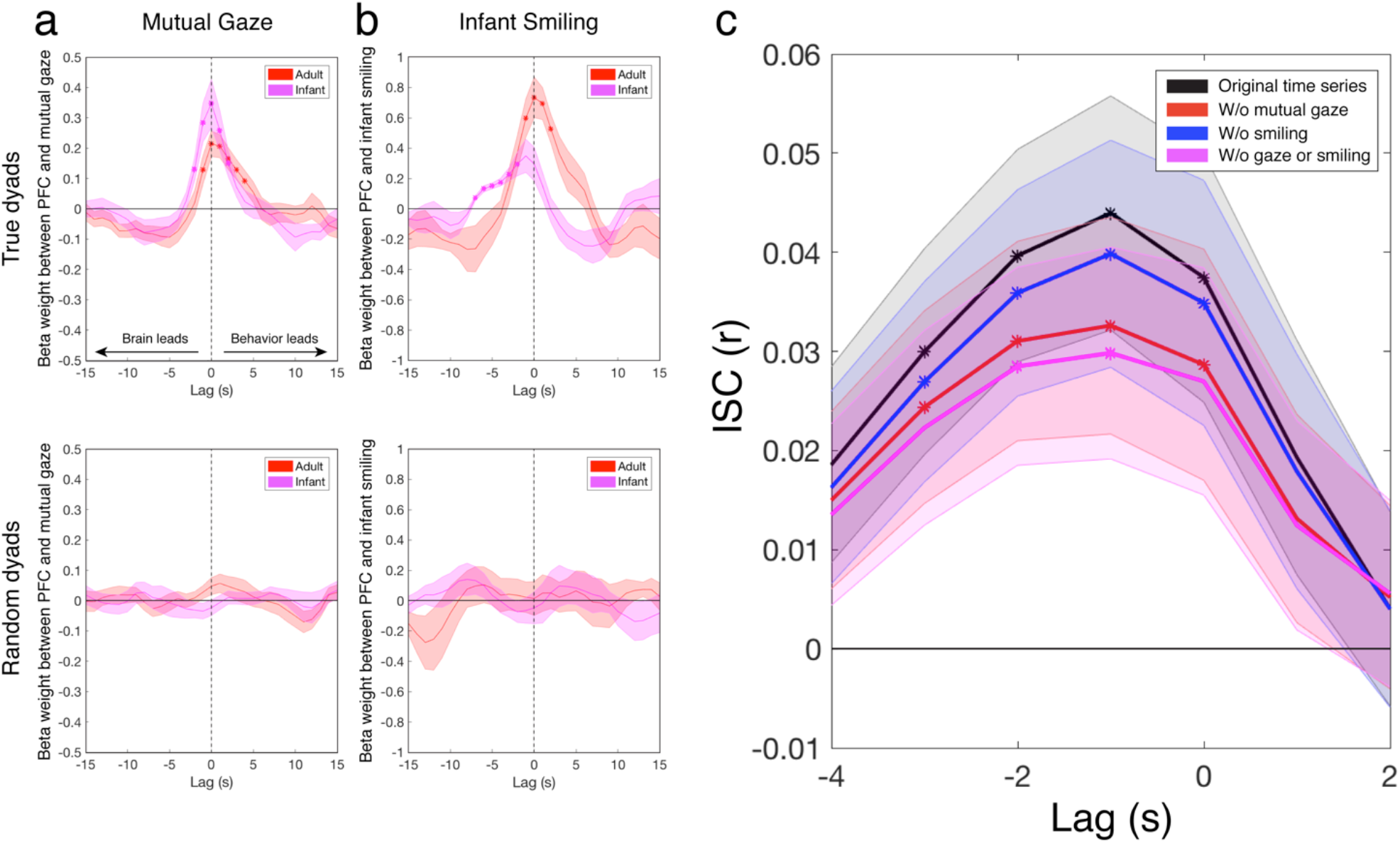
(a, b) Time-lagged regression coefficient between neural responses in the PFC and continuous measures of (a) mutual gaze (*N* = 17) and (b) infant smiling (*N* = 6). Stars indicate time lags at which brain-behavior regression coefficients significantly exceeded 0 (after multiple comparisons correction across lags). Bottom row depicts control results using random brain-behavior assignments. (c) Time-lagged infant-adult coupling (inter-subject correlation) using original PFC time series (black; represents *Together* curve shown in Figure 3A), after regressing out the time course of mutual gaze from the PFC time series (red), after regressing out the time course of infant smiling (blue), and after removing both mutual gaze and smiling (pink). Stars indicate time lags at which ISC significantly exceeded 0 (after FDR correction across lags).

**Figure 5.**
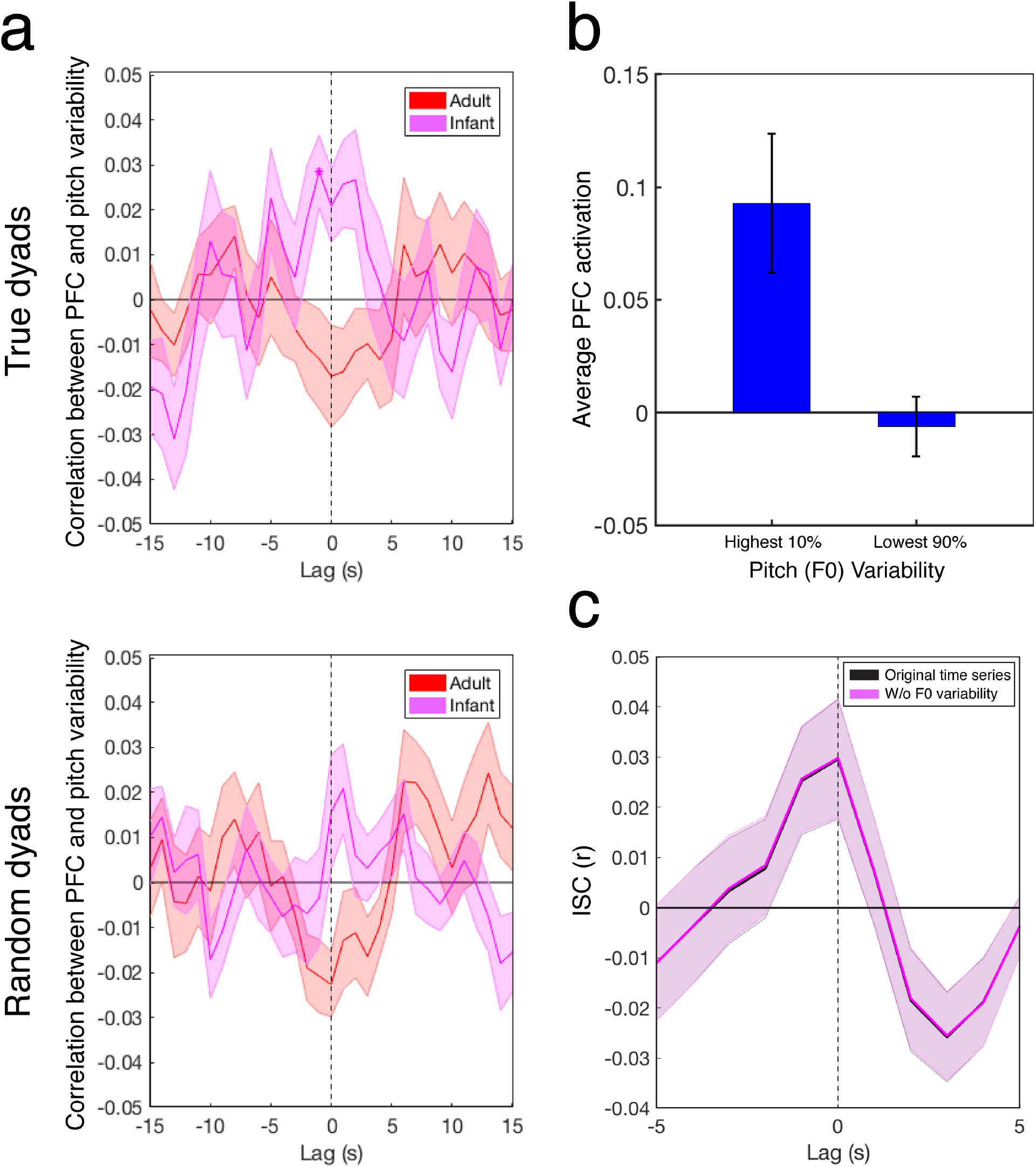
(a) Time-lagged correlation between neural responses in the PFC and a continuous measure of F0 (pitch) variability of the adult’s speech. Stars indicate time lags at which brain-behavior correlations significantly exceeded 0 (after FDR correction across lags). Bottom row depicts control results using random brain-behavior assignments. (b) Infant PFC activation in time windows containing the highest 10% vs. the lower 90% of F0 variability in adult speech for that dyad. (c) Time-lagged infant-adult coupling (inter-subject correlation) using original PFC time series (black) and after regressing out the time course of F0 variability (pink). *N* = 17.

### Infant-adult neural coupling is present only during joint interaction

In the *Together* (but not the *Apart*) condition, we found significant coupling between many PFC channels and some parietal channels of the infant and adult (see Figure 2B). Inter-subject correlation (ISC) between the infant and adult brains was measured during the *Together* and *Apart* conditions, and these actual correlation values were compared to null distributions of correlation values (see Methods). We included all homologous (same channel across brains) and non-homologous (different channels across brains) channel pairings in this analysis. Statistical significance was determined using a permutation procedure based on phase-randomized surrogate data, and multiple comparisons correction was performed across channel pairs using false-discovery rate (FDR, *q* < .05; Benjamini & Hochberg, 1995). These results reveal that direct, real-life communication between an infant and caregiver is reflected in significant inter-subject coupling throughout cortical areas involved in social and narrative processing. The spatial selectivity of this pattern of coupling indicates that the effects are not due to widespread, global arousal throughout the brain.

To further assess infant-adult coupling, we next compared the strength of ISC between the infant and adult brains during the *Together* and *Apart* phases in the PFC channels, which exhibited the most robust coupling in the whole-brain analysis. Each participant had 7 channels covering the PFC. For each dyad, we computed the average ISC across all 49 pairwise combinations of PFC channels (e.g., infant channel 1 vs. adult channel 1, infant channel 1 vs. adult channel 2, etc.). To assess the temporal dynamics of the infant-adult coupling, we shifted the two signals relative to each other in time and performed the above statistical comparison (*Together* vs. *Apart*) at shifts (lags) of -15 seconds (infant leading) to +15 seconds (adult leading), in 1-s increments (Figure 3A).

Significant infant-adult neural coupling was found at lag zero and negative lags of -1, -2, and -3 s (infant leading), but only for the *Together* condition (Figure 3A and Supplementary Fig. 1). Infant-adult neural coupling in the *Apart* condition did not exceed chance at any lag. Similarly, infant-adult neural coupling was significantly greater in the *Together* than the *Apart* condition only at the same set of lags (0, -1, -2, and -3 s; see Supplementary Fig. 1). The pattern of channel combinations that drove these channel-averaged results is visible in Figure 3B (bottom row), which shows *t*-statistics (*Together* – *Apart*) for all 49 pair-wise channel correlations at each lag. These findings suggest that the infant’s PFC may slightly lead the adult’s PFC. For these time-shifted analyses, *p*-values were corrected using false-discovery rate (FDR) across lags (*q* < .05; Benjamini & Hochberg, 1995).

### The PFC tracks dynamic changes in joint eye contact and infant emotion

Infant-adult coupling emerges from continuous feedback between both interlocutors, through eye contact, facial expressions, vocal prosody, and other cues. Which behavioral cues drive the coupling between the infants’ and adult’s brains during the interaction? To start probing this question, we asked an independent rater (whose scores were validated by two additional raters; see Methods) to view and code the video recording of each infant-adult interaction, on a frame-by-frame basis, along two behavioral dimensions (see Methods for details): 1) the presence of mutual gaze (i.e whether the adult and infant were looking directly at each other’s faces), and 2) the signaling of a smile by the infant.

Next, we performed linear regression, using the continuous behavioral ratings to predict the infants’ and adult’s brain responses. We did so to examine whether responses in PFC increased during moments of mutual gaze and moments when the infant smiled. For each dyad, we performed this regression in each channel of the adult and infant. We then averaged the resulting beta weights within our specific region-of-interest (PFC) and computed group-level statistics across dyads (Supplementary Fig. 1). To assess the precise temporal alignment between the neural and behavioral time series, we shifted the two signals relative to each other in time and performed the above steps at lags from -15 seconds (brain leading) to +15 seconds (behavior leading), in 1-s increments (see Methods).

The activity in PFC increased during moments of mutual gaze (i.e joint eye contact between the infant and adult) in both the infants’ and adult’s brains (Figure 4A, top). In both the infants and adult, the increase in activity during mutual gaze was time-locked to the moment of gaze, and this relationship between brain and behavior was significantly above chance for about 1 second before and up to 3 seconds after the initiation of the mutual gaze (see Supplementary Fig. 1).

PFC activity was also modulated by infants’ smiles. Here we noticed a robust time-locked response in the adult’s PFC for the initiation of a smile by the infant (Figure 4B, top). Interestingly, we observed an increase in the infants’ PFC response that began several seconds before the initiation of the smile. A control analysis randomly re-assigning each dyad to the behavioral data of a different dyad (Figure 4A and 4B, bottom) showed no significant brain-behavior relationship for either of the two measures at any lag. This suggests that the PFC’s time-locked tracking of social behavior is not due to similarities in coarse task structure across dyads.

Finally, we assessed to what extent the direct infant-adult B2B coupling (reported in Figure 3 above) was modulated by mutual gaze and the infants’ smiling, by regressing out these two behavioral time courses from the fNIRS signals of each dyad (both infant and adult) before recalculating ISC from lags of -5 (infant leading) to 5 (adult leading) (Figure 4C; black line represents original *Together* data shown in Figure 3A). At the peak of the curve (lag = -1 s), B2B coupling significantly decreased after the removal of the mutual gaze signal (two-tailed *t*-test; *t*(16) = 3.40, *p* < .01), as well as after the decrease of the signal associated with the infants’ smiles (*t*(16) = 2.47, *p* < .05). However, the infant-adult B2B coupling remained significantly greater than chance even after the removal of these two factors (*t*(16) = 2.79, *p* < .05), suggesting that both behavioral signals partially mediate the neural coupling but do not account for it entirely.

### The infant’s PFC continuously tracks the pitch variability of adult speech

Another factor known to facilitate communication with infants is the intonation of infant-directed speech, also known as “motherese”. To assess the influence of infant-directed speech on B2B coupling, we evaluated whether the infant’s PFC tracks the dynamics of the adult’s speech, focusing on F0 variability because of its importance as a cue that influences infant attention (Fernald & Kuhl, 1987) and language processing (Trainor & Desjardins, 2002). To do this, we measured the relationship between F0 variability and the fNIRS signal across time bins in the *Together* condition. We chose a bin size of 200 ms because the syllabic rate of child-directed speech is typically close to 4-5 Hz (Ryan, 2000). In each time bin in which there were recorded fundamental frequency (F0) values, we extracted the standard deviation of F0 of the adult’s voice as well as the average fNIRS activation in that bin. We then computed the correlation between these two measures, averaged across the two most frontal PFC channels (Bonferroni-corrected for all possible pairs of PFC channels, α = .002).

An increase in pitch variability in the caregiver’s speech induced a corresponding increase in PFC response amplitude within the infants. Figure 5A (top) shows that the infant’s brain was significantly correlated with the F0 variability of the adult’s speech at a lag of –1 s (*t*(16) = 3.53, *p* <.001) after FDR correction across lags (from -15 to +15 s, as in the analyses above). The adult’s PFC was not significantly correlated with this measure at any lag, suggesting that the adult’s own pitch variability did not modify the responses in her PFC. Once again, a control analysis randomly re-assigning each dyad to the behavioral data of a different dyad (Figure 5A, bottom) showed no significant brain-behavior relationship at any lag, suggesting that the infant PFC’s tracking of the variability of adult speech is not due to similarities in coarse task structure across dyads.

Next, for each dyad, we found time windows containing the highest 10% of F0 variability for that session and compared infant PFC activation (again, in the first two channels) with activation during all other moments (lower 90% in F0 variability). Moments of particularly high pitch variability had significantly higher infant PFC activation than other moments (Figure 5B; two-tailed *t*-test, *t*(16) = 3.63, *p* < .01).

Finally, we assessed the extent to which direct infant-adult B2B coupling was modulated by the adult speaker’s pitch variability, by regressing its time course from the fNIRS signals of each dyad (both infant and adult) before recalculating ISC at lags of -5 to 5 (Figure 5C). The inclusion of fNIRS data only at time points containing speech (see above) in this analysis resulted in a slightly noisier baseline ISC curve here (black line, Figure 5C) than in Figure 3A (red line), but the shape remained highly consistent. At the peak of the curve (0-lag), we found no significant change in the resulting ISC after removing F0 variability (two-tailed *t*-test; *t*(16) = -1.63, *p* < .12), suggesting that the infant-adult B2B coupling observed in PFC was not mediated by the adult’s speech prosody over time. This finding is consistent with our finding of lack of modulation of the adult’s neural responses in the PFC by the speaker’s F0 variability (Figure 5A, top).

## Discussion

From the beginning of infancy, learning is a dynamic process that hinges on interaction with others. Researchers have uncovered important details about how the behaviors of infants and adults are coupled during natural communication (e.g., Cohn & Tronick, 1988; Marsh, Richardson, & Schmidt, 2009), but very little is known about how their brains dynamically interact during this process. Our research, using an infant-friendly imaging technique, provides the first demonstration of how face-to-face behavioral alignment corresponds with brain-to-brain neural alignment across the developing brain and the mature brain. During coupled interactions, the two brains are dynamically tuned to important social cues – gaze, smiling, and – to some extent – speech prosody. Our findings represent a crucial step toward understanding how infants’ brains begin to extract the most important structure from adults’ input, and how adults’ brains, in turn, represent infants’ emotional feedback as they strive to engage them in everyday communication. This opens new possibilities for understanding the independence versus interdependence of two brains, for evaluating individual differences in early processing and learning, and for uncovering the many dimensions of coupling in infant-caregiver interactions over time.

Our work furthers previous fNIRS research showing heightened mPFC activation in infants in response to direct gaze (Urakawa et al., 2015) and EEG research showing enhanced inter-subject coupling between infants and adults during direct (versus indirect) gaze (Leong et al., 2017). The present study extends these findings by measuring neural coupling during rich, everyday social communication, including interactive playing, singing, and book-reading. Our study demonstrates that the prefrontal cortex of both infants and adults continuously tracks moment-to-moment changes in the interaction, namely in the initiation of eye contact by the infant and adult, and the initiation of smiling by the infant. For each of these behaviors, we uncovered robust, time-locked correlations between neural and behavioral measures. While eye contact and smiling coincide with infant-adult neural coupling, they are not sufficient to account for the entire shared variance across brains. To measure this, we used infants’ and adults’ neural responses to directly search for covaried, shared neural responses across the brains. This method is powerful, as it circumvents the need to rely on an explicit model of the process that drives neural coupling, and as such we were able to detect neural coupling among infant and adult brains above and beyond the coupling that was correlated with gaze and smiling.

The observed interdependent responsiveness of infant and adult brains converges with research showing that humans, from birth, preferentially attend to social and/or communicative information. They prefer speech to structurally similar non-speech sounds (Vouloumanos & Werker, 2004), face-like configurations of dots to unnatural or inverted versions (Farroni et al., 2005), and communicative gestures to non-linguistic pantomime (Krentz & Corina, 2008). This social interest influences early learning. For example, communicative contexts shape what infants remember about objects (Yoon, Johnson, & Csibra, 2008), and infants learn patterns better from communicatively relevant signals, including speech (Marcus, Fernandes, & Johnson, 2007; Rabagliati, Ferguson, & Lew-Williams, in press) or even sine-wave tones that serve a clear communicative function (Ferguson & Lew-Williams, 2016). Moreover, infants’ gaze-following behavior predicts later language outcomes (Brooks & Meltzoff, 2005). This close attention to other people’s communicative cues could shape how the infant brain aligns with adult brains over the course of early development, leading to a highly efficient system for rapidly exchanging information and adopting the norms of a community of speakers. Given the relatively weak neurophysiological response to eye contact in infants at risk for autism spectrum disorder (Elsabbagh et al., 2009), our paradigm invites future research on whether or how brain-to-brain and brain-to-behavior coupling becomes derailed in atypically developing populations.

The findings support transactional development models, which emphasize not only the role of adult input but the child’s role in shaping his own input (Sameroff & Chandler, 1975; Sameroff, 2009; for related ideas, see Smith, Jayaraman, Clerkin, and Yu, 2018). Our results expand on these frameworks and demonstrate a new application of dynamic systems theory (Thelen & Smith, 1996) by showing how infants’ neural activity can also shape the activity in an adult’s brain in highly natural, social interaction. In particular, we found that prefrontal activation in the infant brain precedes and drives similar activation in the adult brain, despite the infant’s very minimal vocalization. This may suggest that the adult is sensitive (likely via a combination of explicit and implicit processes) to subtle behavioral cues from the infant, which in turn dynamically change and modify the adult’s brain responses and behaviors in order to improve neural and behavioral alignment with the developing infant. That is, the brain-to-brain coupling framework suggests that infant and adult brains are engaged in dynamic, closed-loop interactions aimed at maximizing information transfer across brains.

Importantly, and in support of this dynamic view, we measured the alignment between brain and behavior from both sides of the dyadic interaction. The time course of the brain-behavior relationships we report provides preliminary insights into how the brain – in concert with another brain – incorporates live feedback during natural communication. Specifically, in some cases, behavior preceded the neural response, and in others, neural activity preceded behavior. For instance, the PFC of both the infant and adult was significantly coupled to the time course of mutual gaze at lags that skewed slightly positive (Figure 4A, top), implying that behavioral alignment caused neural coupling. This suggests that the brain tracked this social event (which occurs somewhat unpredictably during natural interaction) with a slight delay, due in part to the known hemodynamic lag. However, our smiling-related findings point to a different sequence of events. Although the adult’s PFC was precisely locked to the time course of infant smiling, the infant’s brain preceded his/her own smiling behavior by a few seconds (Figure 4B, top), suggesting that there is a build-up of activation in the PFC (see Habel et al., 2005) even before emotion is outwardly expressed. And the onset of smiling, in turn, preceded the adult’s neural synchronization to the infant.

Even soon after birth, the infant brain is sensitive to the temporal structure of sounds (Telkemeyer et al., 2009). Our finding that the most frontal portion of the PFC tracks changes in F0 variability of adult speech indicates that when the adult uses a more extreme pitch contour to highlight a syllable or word, infant brain activation increases, confirming that the exaggerated sweeps of infant-directed speech provide a salient cue that indeed drives infant attention (Fernald & Kuhl, 1987). This is consistent with the adult mPFC’s tracking of variation in musical features, such as tonality (Janata et al., 2002). In contrast to the infants’ brains, the adult speaker’s PFC responses were not modulated by her own infant-directed speech. In this study, we could only assess infant-adult B2B coupling within and between the PFC and temporal-parietal channels. Future studies might search for coupling between the infant’s PFC and other regions of the adult’s brain more closely related to the production of infant-directed speech.

To conclude, our investigation represents an innovative approach to the study of dynamic social interaction – namely, by tracking the back-and-forth relationships between brains and behaviors during live communication. By studying this process in 1-year-old infants, this approach has the potential to significantly advance models of socially embedded cognition and learning in development. Central to this investigation is that neural coupling is partially mediated by social cues, such as shared gaze and infant smiling, which modify and enhance the dynamics of coupling in both parallel and complementary ways. Our findings highlight the value of taking into account the perspectives of both the developing and mature members of dyads, and prompt further study of the reciprocal, interactive dynamics of brain-to-brain coupling. This will enable a rich understanding of how infants and adults naturally work together to facilitate playful, shared communication.

## Methods

### Participants

Eighteen infants (*M* = 11.3 months, range = 9.8-14.9 months, 9 female), with no history of hearing problems or known developmental delays, participated in the experiment. One experimenter with extensive parenting experience performed the “adult” role in all experimental sessions. Three infants were unable to participate because they refused to wear the fNIRS cap, and 21 additional participants were excluded from statistical analyses because of excessive signal noise or artifacts, typically resulting from the participant grabbing the cap or moving his/her head excessively. One infant was excluded from behavioral analyses due to video malfunction.

### Procedure

We used a dual-brain Shimadzu LabNIRS system to simultaneously record brain activity from the adult and infant in each dyad. The cap measured 57 channels (3-cm in the adult, 2.5-cm in the infants) across the cortex of each participant. These channels covered prefrontal cortex, temporal-parietal junction, and parietal cortex (i.e areas involved in prediction, language processing, and understanding others’ perspectives; Lerner et al., 2011). The locations of these channels were homologous across the infant and adult. Caps were positioned based on known anatomical landmarks according to the 10-20 international system (e.g., center point of the cap was approximately at Cz). Our analyses focused on deoxy-Hb because it is less likely than oxy-Hb to include systemic effects and therefore measures more spatially precise cortical activation (Boas et al., 2014; Hirsch, Zhang, Noah, & Ono, 2017).

All adult-infant dyads participated in two 5-minute conditions. In the *Together* condition, the adult experimenter engaged directly with the child by playing with a consistent set of toys, singing nursery rhymes, and reading *Goodnight Moon*. The child sat on his/her parent’s lap, and the parent was told to keep the child comfortable but not to communicate with the child in any way. In the control (*Apart*) condition, the experimenter turned 90 degrees away from the child and told a story to an adult experimenter using adult-directed speech, while the child interacted quietly with his/her parent. The order of the two conditions was counterbalanced across participants. This comparison allowed us to test whether coupling was stronger when the adult and child directly communicated with each other than when they were engaged in a similarly communicative task, but not with each other.

### Pre-processing and analysis

We removed motion artifacts using moving standard deviation and spline interpolation (Scholkmann, Spichtig, Muehlemann, & Wolf, 2010), and also low-pass (0.5 Hz) and high-pass (0.02 Hz) filtered the signal to remove physiological noise and drift, respectively. To compute inter-subject correlation, we computed a Pearson correlation between one channel in the adult and a channel in the infant (Figure 1).

Because of the presence of long-range temporal autocorrelation in fNIRS time series, we estimated the statistical likelihood of each observed correlation using a bootstrapped permutation procedure based on surrogate data generated using phase randomization (see Simony et al., 2016), which preserves the mean and autocorrelation of the original signal but randomizes the phases after applying a Fast Fourier Transform. For each of 3,249 channel combinations (57 infant x 57 adult), we computed a group-averaged Pearson correlation coefficient based on phase-scrambled time series in the adult and infant of each dyad. We performed this procedure 20,000 times to yield a null distribution of ISCs for each channel combination and computed a *p*-value by calculating the proportion of null values that exceeded the true ISC value for that channel combination. *P*-values of 0 (in which the entire null distribution fell below the true ISC) were set to 1 ÷ (the number of bootstrapped samples). To correct for multiple comparisons, we applied the false-discovery rate (FDR) procedure (Benjamini & Hochberg, 1995), with a *q*-value of .05. As shown in Figure 2B, this resulted in eleven significant channel pairs (primarily in the PFC) in the *Together* condition and no significant channel pairs in the *Apart* condition.

During the *Together* condition, we continuously recorded video and audio of the dyadic interaction. Later, a research assistant coded every 500 ms whether or not there was mutual gaze (joint eye contact) between the adult and infant, and whether or not the infant was smiling. We did not code adult smiling because the adult experimenter smiled >95% of the time; the nearly constant presence of adult smiling left minimal opportunities for the infant to react to individual instances of smiling. Two additional research assistants each coded mutual gaze and smiling in subsets of the data (two-thirds of the datasets each), and the scores were highly reliable across the three raters (Cronbach’s α = .70, averaged across the two behavioral measures; Cronbach, 1951). After filtering out background noise from the audio recordings in Adobe Audition, we analyzed continuous fundamental frequency (F0) using Praat. Only six infants displayed any smiling throughout the videos, and this subset was used in the smiling-related analyses.

To perform brain-behavior comparisons, we first down-sampled the fNIRS time series in each channel to match the behavioral data (2 Hz for mutual gaze and smiling, 5 Hz for F0 variability) before computing correlations.

## Acknowledgments

This work was supported by the Princeton University C. V. Starr Fellowship (E.A.P.), the Eric and Wendy Schmidt Transformative Technology Award (E.A.P., U.H., C.L.W.), NIH 1DP1HD091948 (U.H.), and NIH HD079779 (C.L.W.). We thank Ariella Cohen for assistance with data collection and behavioral coding, Alice Wang and Mia Sullivan for assistance with behavioral coding, and Eva Fourakis, Carolyn Mazzei, and other members of the Princeton Baby Lab for help with participant recruitment.

---

**Supplementary Figure 1.**
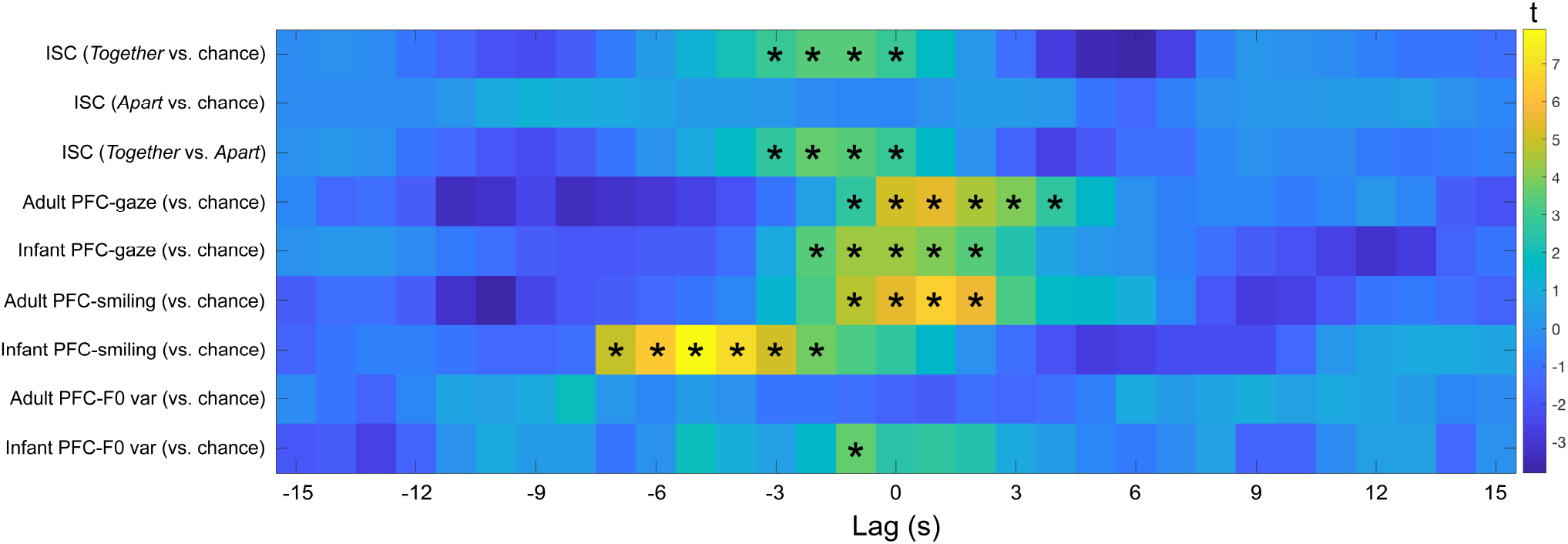
Statistical results (t-statistics) for inter-subject correlation analyses (rows 1-3), brain-behavior regression analyses (mutual gaze, infant smiling; rows 4-7), and brain-behavior correlation analyses (F0 variability of adult speech; rows 8-9). Stars indicate all t-values that reached significance after multiple comparisons correction across lags (*q* < .05 for all analyses; Benjamini & Hochberg, 1995).

